# Tools and techniques for computational reproducibility

**DOI:** 10.1101/022707

**Authors:** Stephen R. Piccolo, Michael B. Frampton

**Affiliations:** Department of Biology, Brigham Young University, Provo, UT, USA; Department of Computer Science, Brigham Young University, Provo, UT, USA

## Abstract

When reporting research findings, scientists document the steps they followed so that others can verify and build upon the research. When those steps have been described in sufficient detail that others can retrace the steps and obtain similar results, the research is said to be reproducible. Computers play a vital role in many research disciplines and present both opportunities and challenges for reproducibility. Computers can be programmed to execute analysis tasks, and those programs can be repeated and shared with others. Due to the deterministic nature of most computer programs, the same analysis tasks, applied to the same data, will often produce the same outputs. However, in practice, computational findings often cannot be reproduced due to complexities in how software is packaged, installed, and executed—and due to limitations in how scientists document analysis steps. Many tools and techniques are available to help overcome these challenges. Here we describe seven such strategies. With a broad scientific audience in mind, we describe strengths and limitations of each approach, as well as circumstances under which each might be applied. No single strategy is sufficient for every scenario; thus we emphasize that it is often useful to combine approaches.

## Introduction

When reporting research, scientists document the steps they followed to obtain their results. If the description is comprehensive enough that they and others can repeat the procedures and obtain semantically consistent results, the findings are considered to be “reproducible”^1–6^. Reproducible research forms the basic building blocks of science, insofar as it allows researchers to verify and build on each other's work with confidence.

Computers play an increasingly important role in many scientific disciplines^7–10^. For example, in the United Kingdom, 92% of academic scientists use some type of software in their research, and 69% of scientists say their research is feasible only with software tools^11^. Thus efforts to increase scientific reproducibility should consider the ubiquity of computers in research.

Computers present both opportunities and challenges for scientific reproducibility. On one hand, due to the deterministic nature of most computer programs, many computational analyses can be performed such that others can obtain exactly identical results when applied to the same input data^12^. Accordingly, computational research can be held to a high reproducibility standard. On the other hand, even when no technical barrier prevents reproducibility, scientists often cannot reproduce computational findings due to complexities in how software is packaged, installed, and executed—and due to limitations in how scientists document these steps^13^. This problem is acute in many disciplines, including genomics, signal processing, and ecological modeling^14–16^, where data sets are large and computational tools are evolving rapidly. However, the same problem can affect any scientific discipline that requires computers for research. Seemingly minor differences in computational approaches can have major influences on analytical outputs^12,17–21^, and the effects of these differences may exceed those that result from experimental factors^22^.

Journal editors, funding agencies, governmental institutions, and individual scientists have increasingly made calls for the scientific community to embrace practices that support computational reproducibility^23–30^. This movement has been motivated, in part, by scientists' failed efforts to reproduce previously published analyses. For example, Ioannidis, et al. evaluated 18 published research studies that used computational methods to evaluate gene-expression data but were able to reproduce only 2 of those studies^31^. In many cases, a failure to share the study's data was the culprit; however, incomplete descriptions of software-based analyses were also common. Nekrutenko and Taylor examined 50 papers that analyzed next-generation sequencing data and observed that fewer than half provided any details about software versions or parameters^32^. Recreating analyses that lack such details can require hundreds of hours of effort^33^ and may be impossible, even after consulting the original authors. Failure to reproduce research may also lead to careerist effects, including retractions^34^.

Noting such concerns, some journals have emphasized the value of placing computer source code in open-access repositories, such as *GitHub* (https://github.com) or *BitBucket* (https://bitbucket.org). In addition, journals have extended requirements for “Methods" sections, now asking researchers to provide detailed descriptions of 1) how to install software and its dependencies and 2) what parameters and data-preprocessing steps are used in analyses^10,23^. A recent Institute of Medicine report emphasized that, in addition to computer code and research data, fully specified computational procedures should be made available to the scientific community^24^. They elaborated that such procedures should include “all of the steps of computational analysis" and that “all aspects of the analysis need to be transparently reported”^24^. Such policies represent important progress. However, it is ultimately the responsibility of individual scientists to ensure that others can verify and build upon their analyses.

Describing a computational analysis sufficiently—such that others can reexecute it, validate it, and refine it—requires more than simply stating what software was used, what commands were executed, and where to find the source code^13,26,35–37^. Software is executed within the context of an operating system (for example, *Windows, Mac OS*, or *Linux*), which enables the software to interface with computer hardware (Figure 1). In addition, most software relies on a hierarchy of software dependencies, which perform complementary functions and must be installed alongside the main software tool. One version of a given software tool or dependency may behave differently or have a different interface than another version of the same software. In addition, most analytical software offers a range of parameters (or settings) that the user can specify. If any of these variables differs from what the original experimenter used, the software may not execute properly or analytical outputs may differ considerably from what the original experimenter observed.

**Figure 1.**
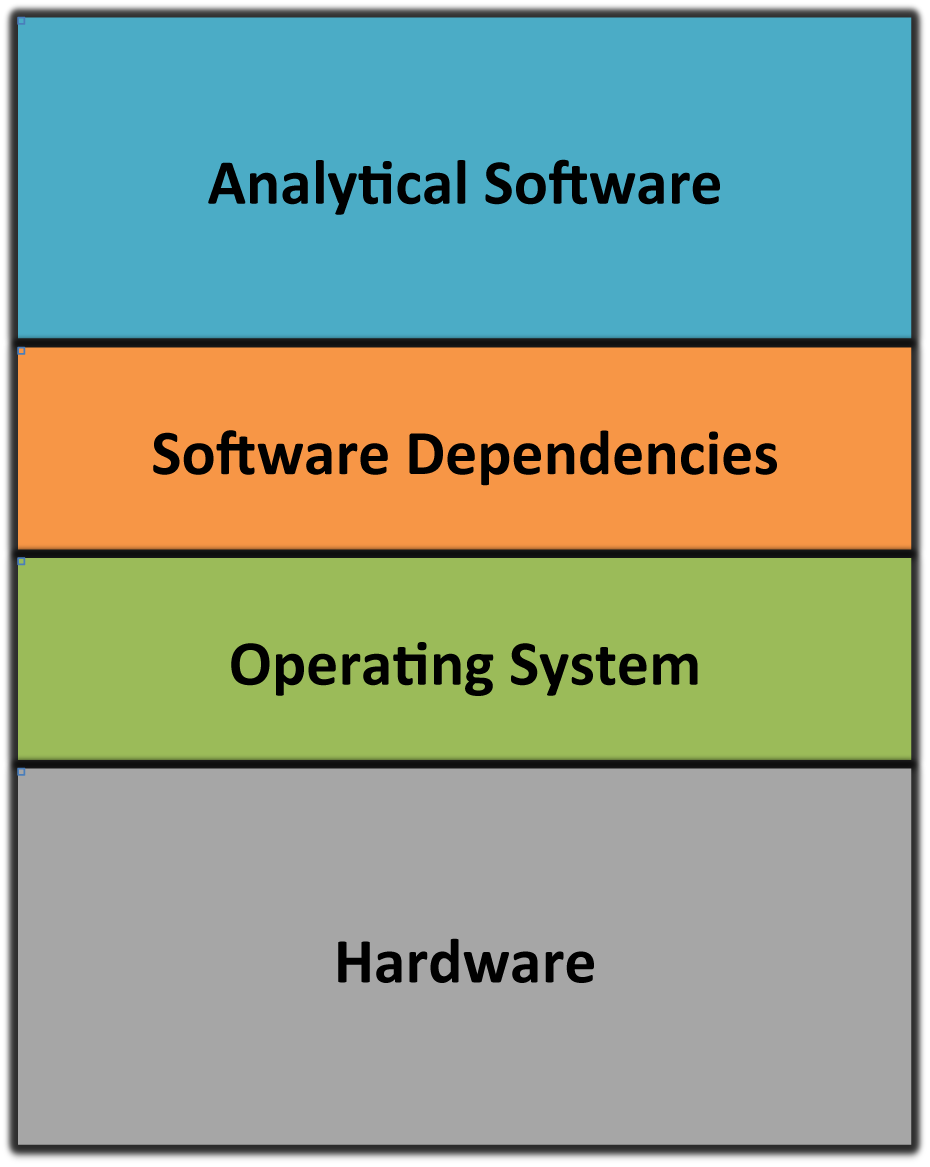
Basic computer architecture. Computer hardware consists of hardware devices, including central processing units, hard drives, random access memory, keyboard, mouse, etc. Operating systems enable software to interface with hardware; popular operating-system families are *Windows, Mac OS*, and *Linux*. Users interact with computers via software interfaces. In scientific computing, software enables users to execute algorithms, analyze data, generate graphics, etc. To execute properly, most software tools depend on specific versions of software dependencies, which must be installed on the same operating system.

Scientists can use various tools and techniques to overcome these challenges and to increase the likelihood that their computational analyses will be reproducible. These techniques range in complexity from simple (e.g., providing written documentation) to advanced (e.g., providing a “virtual" environment that includes an operating system and all software necessary to execute the analysis). This review describes seven strategies across this spectrum. We describe strengths and limitations of each approach, as well as circumstances under which each might be applied. No single strategy will be sufficient for every scenario; therefore, in many cases, it will be most practical to combine multiple approaches. This review focuses primarily on the computational aspects of reproducibility. The related topics of empirical reproducibility, statistical reproducibility, and data sharing have been described elsewhere^38–44^. We believe that with greater awareness and understanding of computational-reproducibility techniques, scientists—including those with limited computational experience—will be more apt to perform computational research in a reproducible manner.

## Narrative descriptions are a simple but valuable way to support computational reproducibility

The most fundamental strategy for enabling others to reproduce a computational analysis is to provide a detailed, written description of the process. For example, when reporting computational results in a research article, authors customarily provide a narrative that describes the software they used and the analytical steps they followed. Such narratives can be invaluable in enabling others to evaluate the scientific approach and to reproduce the findings. In many situations—for example, when software execution requires user interaction or when proprietary software is used—narratives are the only feasible option for documenting such steps. However, even when a computational analysis uses open-source software and can be fully automated, narratives help others understand how to reexecute an analysis.

Although most research articles that use computational methods provide some type of narrative, these descriptions often lack sufficient detail to enable others to retrace those steps ^31,32^. Narrative descriptions should indicate the operating systemfs), software dependencies, and analytical software that were used and how to obtain them. In addition, narratives should indicate the exact software versions used, the order in which they were executed, and all non-default parameters that were specified. Such descriptions should account for the fact that computer configurations differ vastly, even for computers that use the same operating system. Because it can be difficult for scientists to remember such details after the fact, it is best to record this information throughout the research process, rather than at the time of manuscript preparation^8^.

The following sections describe techniques for automating computational analyses. These techniques can diminish the need for scientists to write narratives. However, because it is often impractical to automate all computational steps, we expect that, for the foreseeable future, narratives will play a vital role in enabling computational reproducibility.

## Custom scripts and code can automate a research analysis

Scientific software can often be executed in an automated manner via text-based commands. Using such commands—via a *command-line interface*—scientists can indicate which software program(s) should be executed and which parameter(s) should be used. When multiple commands must be executed, they can be compiled into *scripts*, which specify the order in which the commands should be executed (Figure 2). In many cases, scripts also include commands for installing and configuring software. Such scripts serve as valuable documentation not only for individuals who wish to reexecute the analysis but also for the researcher who performed the original analysis^45^. In these cases, no amount of narrative is an adequate substitute for providing the actual commands that were used.

**Figure 2.**
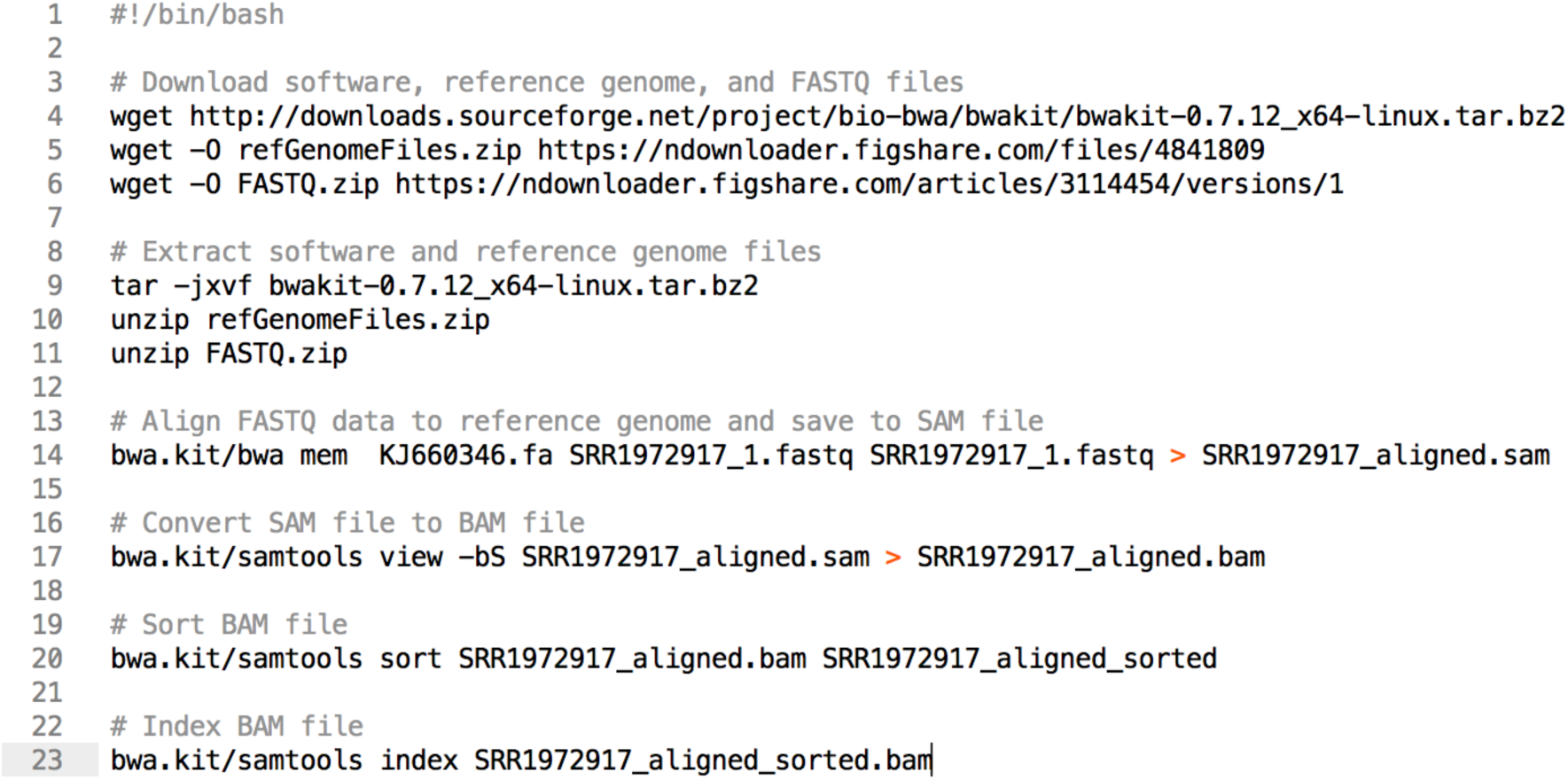
Example of a command-line script. This script can be used to align DNA sequence data to a reference genome. First it downloads software and data files necessary for the analysis. Then it extracts (“unzips”) these files, aligns the data to a reference genome for Ebolavirus. Finally, it converts, sorts, and indexes the aligned data.

When writing command-line scripts, it is essential to explicitly document any software dependencies and input data that are required for each step in the analysis. The *Make* utility^46^ provides one way to specify such requirements^35^. Before any command is executed, *Make* verifies that each documented dependency is available. Accordingly, researchers can use *Make* files (scripts) to specify a full hierarchy of operating-system components and dependent software that must be present to perform the analysis (Figure 3). In addition, *Make* can be configured to automatically identify any commands that can be executed in parallel, potentially reducing the amount of time required to execute the analysis. Although *Make* was designed originally for UNIX-based operating systems (such as *Mac OS* or *Linux*), similar utilities have since been developed for *Windows* operating systems^47^. Box 1 lists various utilities that can be used to automate software execution.

**Figure 3.**
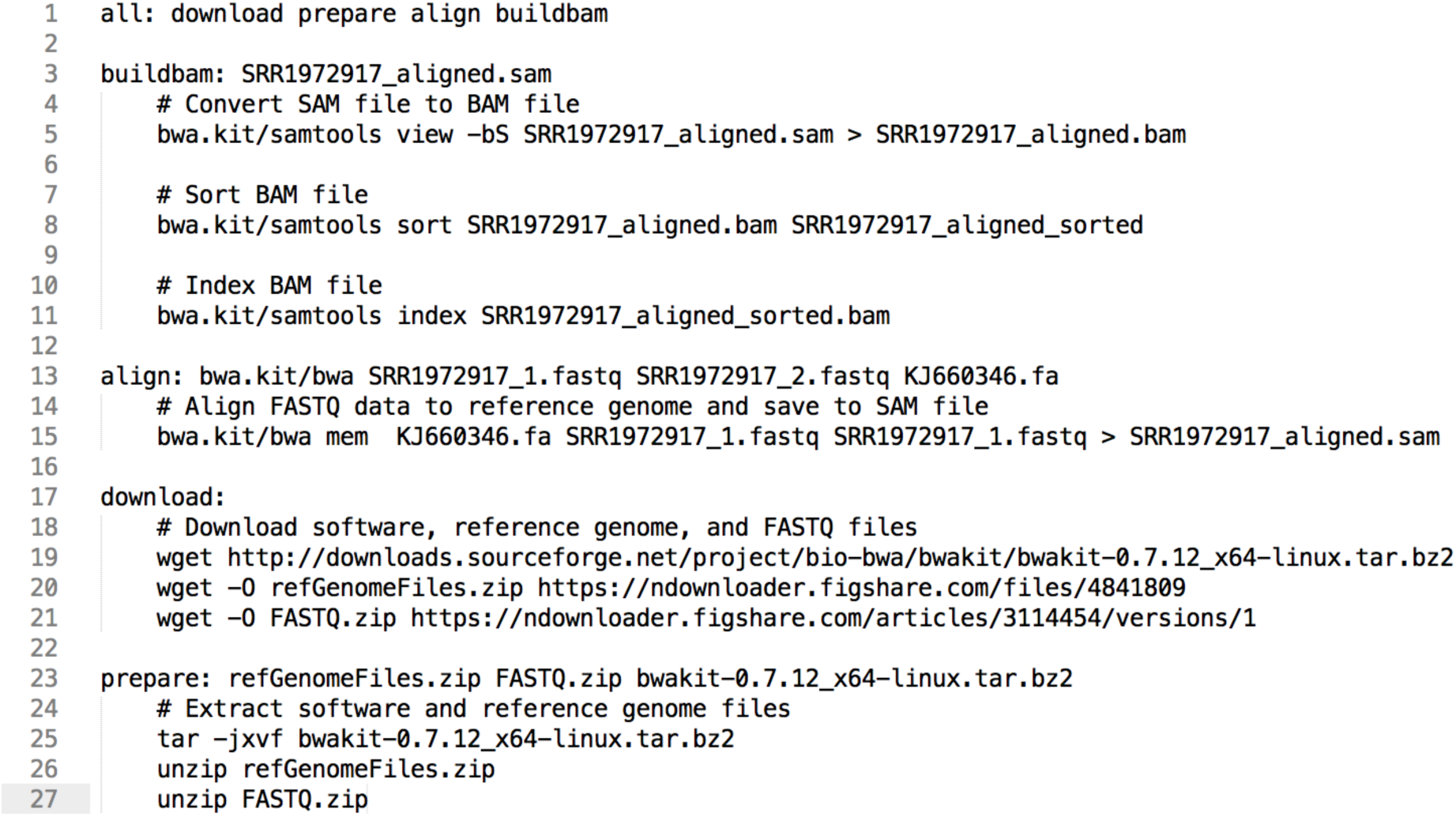
Example of a *Make* file. This file performs the same function as the command-line script shown in Figure 2, except that it is formatted for the *Make* utility. Accordingly, it is structured so that specific tasks must be executed before other tasks, in a hierarchical structure. For example, the “download” target must be completed before the downloaded files can be prepared (“prepare” target). The “all” target specifies all of the targets that must be executed, in order.

#### Box 1 Utilities that can be used to automate software execution

- *GNU Make*^46^ and *Make for Windows*^47^: Tools for building software from source files and for ensuring that the software's dependencies are met.
- *Snakemake*^48^ = An extension of Make that provides a more flexible syntax and makes it easier to execute tasks in parallel.
- *BPipe*^49^ = A tool that provides a flexible syntax for users to specify commands to be executed; it maintains an audit trail of all commands that have been executed.
- *GNU Parallel*^50^ = A tool for executing commands in parallel across one or more computers.
- *Makeflow*^51^ = A tool that can execute commands simultaneously on various types of computer architectures, including computer clusters and cloud environments.
- *SCONS*^52^ = An alternative to *GNU Make* that enables users to customize the process of building and executing software using scripts written in the Python programming language.
- *CMAKE*^53^ = A tool that enables users to execute *Make* scripts more easily on multiple operating systems.

In addition to creating scripts that execute existing software, many researchers also create new software by writing computer code in a programming language such as Python, `C++`, Java, or R. Such code may perform relatively simple tasks, such as reformatting data files or invoking third-party software. In other cases, computer code may constitute a manuscript's key intellectual contribution.

Whether analysis steps are encoded in scripts or as computer code, scientists can support reproducibility by publishing these artefacts alongside research papers. By doing so, the authors enable readers to evaluate the analytical approach in full detail and to extend the analysis more readily^54^. Although scripts and code may be included alongside a manuscript as supplementary material, a better alternative is to store them in a *version-control system* (VCS)^8,9,45^ and to share these repositories via Web-based services like *GitHub* (https://github.com) or *Bitbucket* (https://bitbucket.org). With such a VCS repository, scientists can track different versions of scripts and code that have been developed as the research project evolved. In addition, outside observers can see the full version history, contribute revisions to the code, and reuse the code for their own purposes^55^. When submitting a manuscript, the authors may “tag” a specific version of the repository that was used for the final analysis described in the manuscript.

## Software frameworks enable easier handling of software dependencies

Virtually all computer scripts and code rely on external software dependencies and operating-system components. For example, suppose that a research study required a scientist to apply *Student’s t-test*. Rather than write code that implements this statistical test, the scientist would likely find an existing software *library* that implements the test and then invoke that library from her code. A considerable amount of time can be saved with this approach, and a wide range of software libraries are freely available. However, software libraries change frequently—invoking the wrong version of a library may result in an error or an unexpected output. Thus to enable others to reproduce an analysis, it is critical to indicate which dependencies (and versions thereof) must be installed.

One way to address this challenge is to build on a preexisting software framework. Such frameworks make it easier to access software libraries that are commonly used to perform specific types of analysis task. Typically, such frameworks also make it easier to download and install software dependencies and ensure that the versions of software libraries and their dependencies are compatible with each other. For example, *Bioconductor*^56^, created for the *R* statistical programming language^57^, is a popular framework that contains hundreds of software packages for analyzing biological data^56^. The *Bioconductor* framework facilitates versioning, documenting, and distributing code. Once a software library has been incorporated into *Bioconductor*, other researchers can find, download, install, and configure it on most operating systems with relative ease. In addition, *Bioconductor* installs software dependencies automatically. These features ease the process of performing an analysis and can help with reproducibility. Various software frameworks exist for other scientific disciplines^58–63^. General-purpose tools for managing software dependencies also exist (e.g., *Apache Ivy*^64^ and Puppet^65^).

To best support reproducibility, software frameworks should make it easy for scientists to download and install previous versions of a software tool as well as previous versions of dependencies. Such a design would enable other scientists to reproduce analyses that were conducted with previous versions of a software framework. In the case of *Bioconductor*, considerable extra work may be required to install specific versions of *Bioconductor* software and their dependencies. To overcome such limitations, scientists may use a software container or virtual machine fsee below) to package the specific versions they used in an analysis. Alternatively, they may use third-party solutions such as the aRchive project (http://bioarchive.github.io).

## Literate programming combines narratives directly with code

Although narratives, scripts, and computer code support reproducibility individually, additional value can be gained from combining these entities. Even though a researcher may provide computer code alongside a research paper, other scientists may have difficulty interpreting how the code accomplishes specific tasks. A longstanding way to address this problem is via code comments, which are human-readable annotations interspersed throughout computer code. However, code comments and other types of documentation often become outdated as code evolves throughout the analysis process^66^. One way to overcome this problem is to use a technique called *literate programming*^67^. With this approach, the scientist writes a narrative of the scientific analysis and intermingles code directly within the narrative. As the code is executed, a document is generated that includes the code, narratives, and any outputs (e.g., figures, tables) that the code produces. Accordingly, literate programming helps ensure that readers understand exactly how a particular research result was obtained. In addition, this approach motivates the scientist to keep the target audience in mind when performing a computational analysis, rather than simply to write code that a computer can parse^67^. Consequently, by reducing barriers of understanding among scientists, literate programming can help to engender greater trust in computational findings.

One popular literate-programming tool is *Jupyter*^*68*^. Using its Web-based interface, scientists can create interactive notebooks that combine code, data, mathematical equations, plots, and rich media^69^. Originally known as *IPython* and previously designed exclusively for the *Python* programming language, *Jupyter*^*70*^ now makes it possible to execute code in many different programming languages. Such functionality may be important to scientists who prefer to combine the strengths of different programming languages.

*knitr*^*71*^ has also gained considerable popularity as a literate-programming tool. It is written in the *R* programming language and thus can be integrated seamlessly with the array of statistical and plotting tools available in that environment. However, like *Jupyter, knitr* can execute code written in multiple programming languages. Commonly, *knitr* is applied to documents that have been authored using *RStudio*^*72*^, an open-source tool with advanced editing and package-management features.

*Jupyter* notebooks and *knitr* reports can be saved in various output formats, including HTML and PDF (see examples in Figures 4–5). Increasingly, scientists include such documents with journal manuscripts as supplementary material, enabling others to repeat analysis steps and recreate manuscript figures^73–76^.

**Figure 4.**
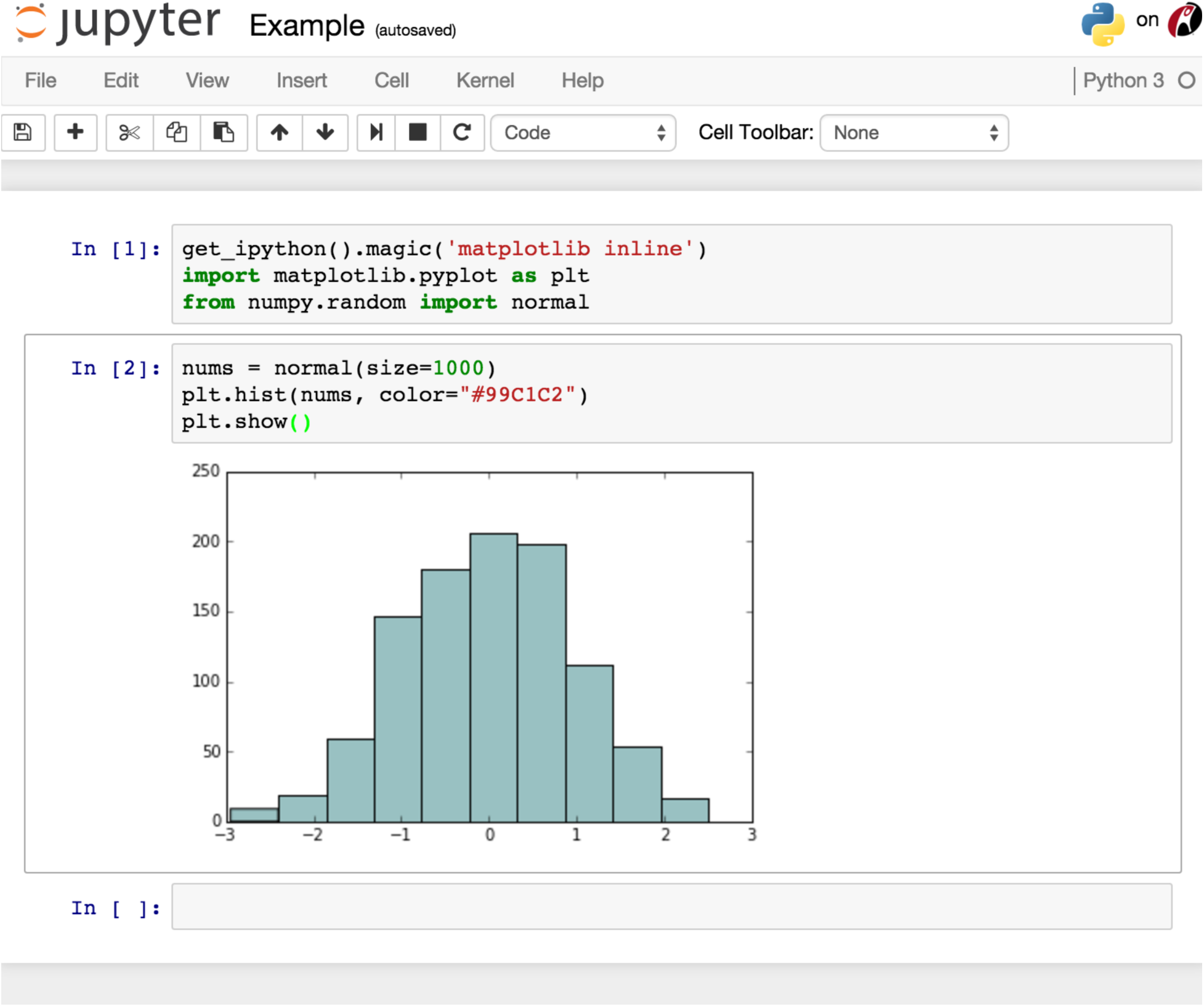
Figure 4: Example of a *Jupyter* notebook. This example contains code (in the Python programming language) for generating random numbers and plotting them in a graph within a *Jupyter* notebook. Importantly, the code and output object (graph) are contained within the same document.

**Figure 5.**
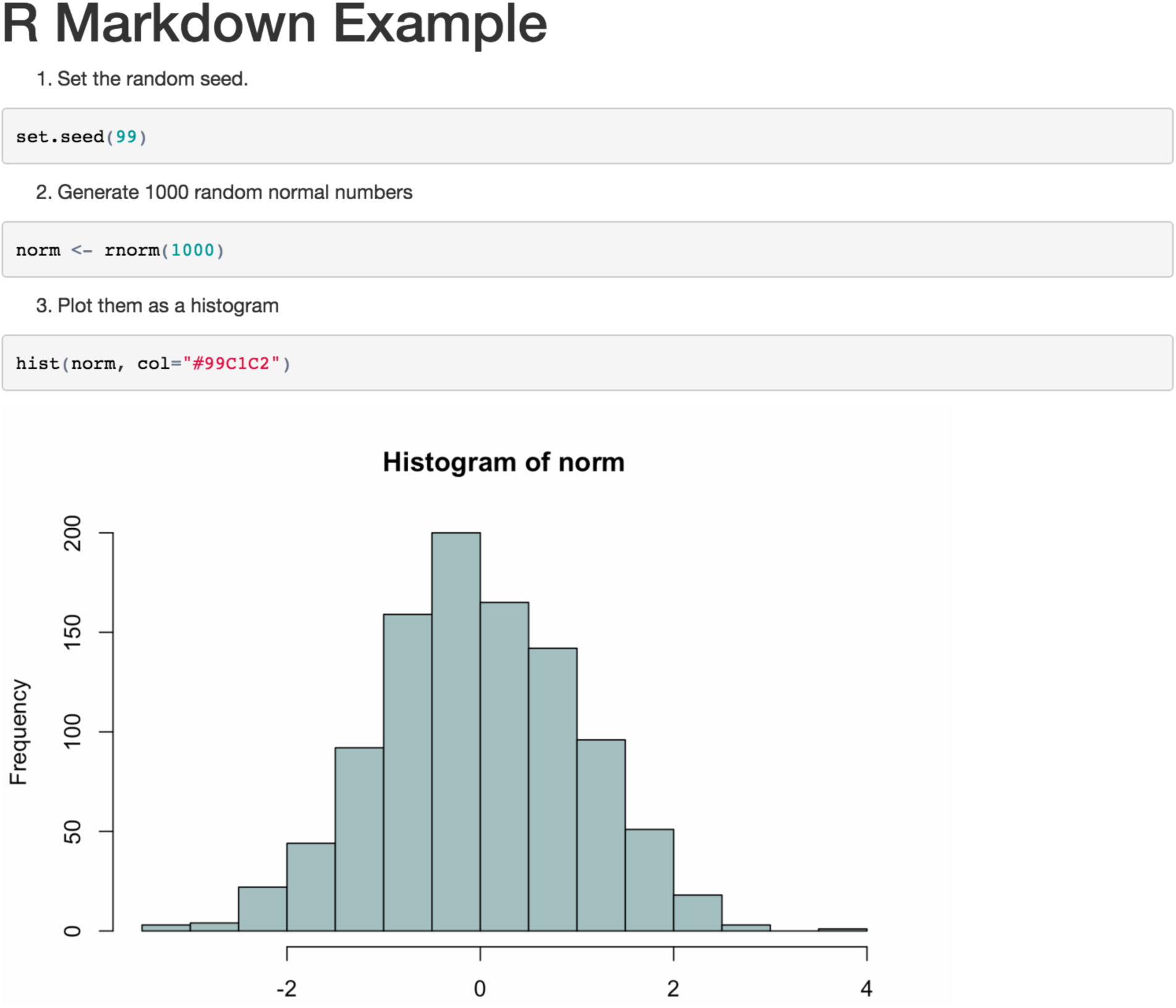
Example of a document that has been created using *knitr*. This example contains code (in the R language) for generating random numbers and plotting them in a graph. The *knitr* tool was used to generate the document, which combines the code and the output object (figure).

Scientists typically use literate-programming tools for data analysis tasks that can be executed in a modest amount of time (e.g., minutes or hours). It is possible to execute *Jupyter* or *knitr* at the command line; thus longer-running tasks can be executed on high-performance computers. However, this approach runs counter to the interactive nature of notebooks and require additional technical expertise to configure and execute the notebooks.

Literate-programming notebooks are suitable for research analyses that require a modest amount of computer code. For analyses that require larger amounts of code, more advanced programming environments may be more suitable—perhaps in combination with a “literate documentation” tool such as Dexy (http://www.dexy.it).

## Workflow-management systems enable software execution via a graphical user interface

Writing computer scripts and code may seem daunting to many researchers. Although various courses and tutorials are helping to make this task less formidable^77–80^, many scientists use “workflow management systems" to facilitate the process of executing scientific software^81^. Typically managed via a graphical user interface, workflow management systems enable scientists to upload data and process it using existing tools. For multistep analyses, the output from one tool can be used as input to additional tools, potentially resulting in a series of commands known as a workflow.

*Galaxy*^*82, 83*^ has gained considerable popularity within the bioinformatics community—especially for performing next-generation sequencing analysis. As users construct workflows, *Galaxy* provides descriptions of how software parameters should be used, examples of how input files should be formatted, and links to relevant discussion forums. To help with processing large data sets and computationally complex algorithms, *Galaxy* also provides an option to execute workflows on cloud-computing services^84^. In addition, researchers can share workflows with each other^85^; this feature has enabled the *Galaxy* team to build a community that helps to encourage reproducibility, define best practices, and reduce the time required for novices to get started.

Various other workflow systems are freely available to the research community (see Box 2). For example, *VisTrails* is used by researchers from many disciplines, including climate science, microbial ecology, and quantum mechanics^86^. It enables scientists to design workflows visually, connecting data inputs with analytical modules and the resulting outputs. In addition, *VisTrails* tracks a full history of how each workflow was created. This capability, referred to as retrospective provenance, makes it possible for others not only to reproduce the final version of an analysis but also to examine previous incarnations of the workflow and examine how each change influenced the analytical outputs^87^.

#### Box 2 Workflow management tools freely available to the research community

- Galaxy^82,83^ - https://usegalaxy.org
- VisTrails^86^ - http://www.vistrails.org
- Kepler^88^ - https://kepler-project.org
- iPlant Collaborative^89^ - http://www.iplantcollaborative.org
- GenePattern^90,91^ - http://www.broadinstitute.org/cancer/software/genepattern
- Taverna^92^ - http://www.taverna.org.uk
- LONI Pipeline^93^ - http://pipeline.bmap.ucla.edu

Although workflow-management systems offer many advantages, users must accept tradeoffs. For example, although the teams that develop these tools often provide public servers where users can execute workflows, many scientists share these limited resources, so the public servers may not have adequate computational power or storage space to execute large-scale analyses in a timely manner. As an alternative, many scientists install these systems on their own computers; however, configuring and supporting them requires time and expertise. In addition, if a workflow tool does not yet provide a module to support a given analysis, the scientist must create a new module to support it. This task constitutes additional overhead; however, utilities such as the *Galaxy Tool Shed*^*94*^ are helping to facilitate this process.

## Virtual machines encapsulate an entire operating system and software dependencies

Whether an analysis is executed at the command line, within a literate-programming notebook, or via a workflow-management system, an operating system and relevant software dependencies must be installed before the analysis can be performed. The process of identifying, installing, and configuring such dependencies consumes a considerable amount of scientists' time. Different operating systems (and versions thereof) may require different installation and configuration steps. Furthermore, earlier versions of software dependencies, which may currently be installed on a given computer, may be incompatible with—or produce different outputs than—newer versions.

One solution is to use virtual machines, which can encapsulate an entire operating system and all software, scripts, code, and data necessary to execute a computational analysis^95,96^ (Figure 6). Using virtualization software—such as *VirtualBox* or *VMWare* (see Box 3)—a virtual machine can be executed on practically any desktop, laptop, or server, irrespective of the main (“host”) operating system on the computer. For example, even though a scientist's computer may be running a *Windows* operating system, the scientist may perform an analysis on a *Linux* operating system that is running concurrently—within a virtual machine—on the same computer. The scientist has full control over the virtual (“guest”) operating system and thus can install software and modify configuration settings as necessary. In addition, a virtual machine can be constrained to use specific amounts of computational resources (e.g., computer memory, processing power), thus enabling system administrators to ensure that multiple virtual machines can be executed simultaneously on the same computer without impacting each other's performance. After executing an analysis, the scientist can export the entire virtual machine to a single, binary file. Other scientists can then use this file to reconstitute the same computational environment that was used for the original analysis. With a few exceptions (see Discussion), these scientists will obtain exactly the same results that the original scientist obtained. This process provides the added benefits that 1) the scientist must only document the installation and configuration steps for a single operating system, 2) other scientists need only install the virtualization software and not individual software components, and 3) analyses can be reexecuted indefinitely, so long as the virtualization software remains compatible with current computer systems^97^. Also useful, a team of scientists can employ virtual machines to ensure that each team member has the same computational environment, even though the team members may have different configurations on their host operating systems.

**Figure 6.**
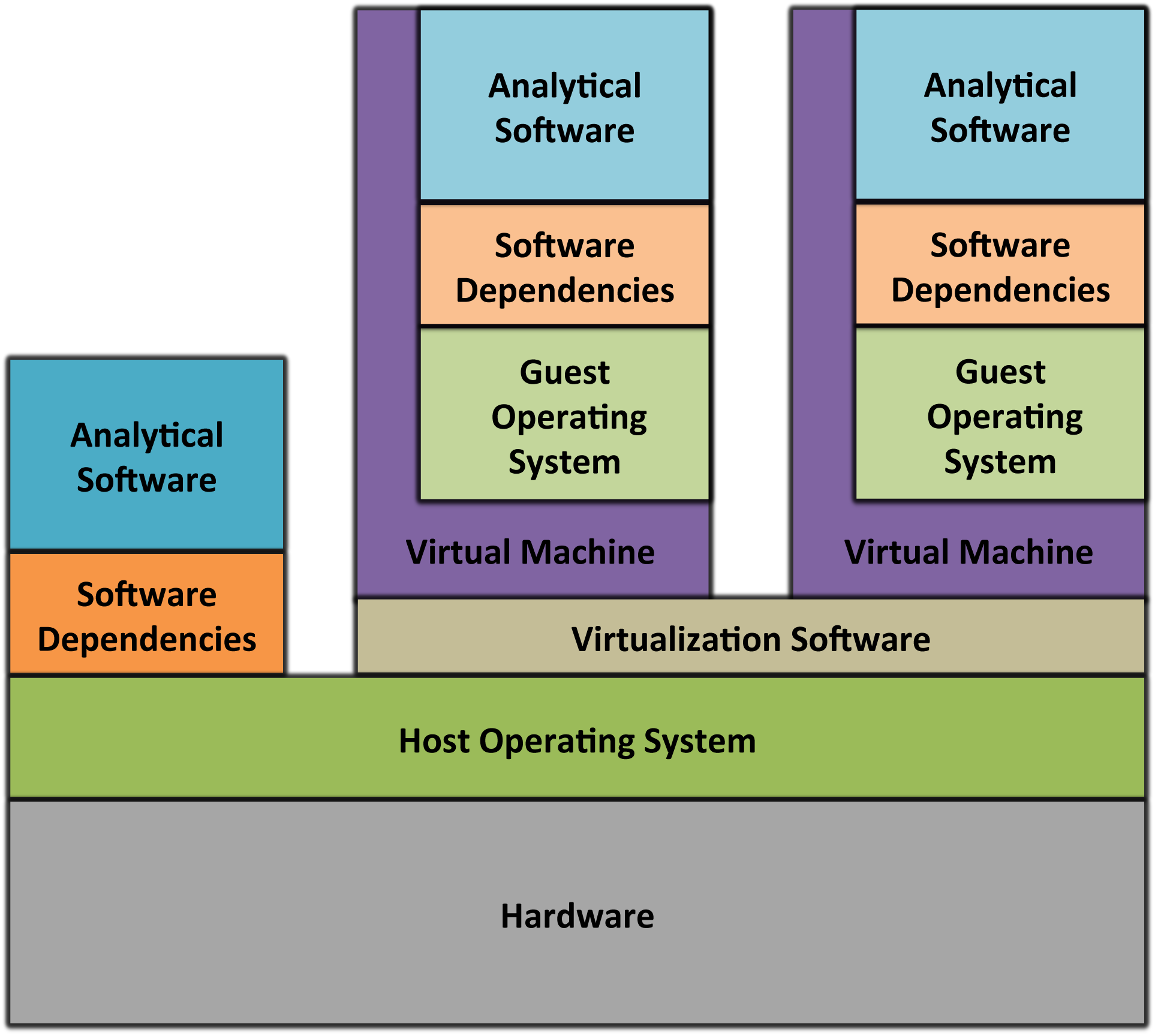
Architecture of virtual machines. Virtual machines encapsulate analytical software and dependencies within a “guest" operating system, which may be different than the main (“host”) operating system. A virtual machine executes in the context of virtualization software, which executes alongside whatever other software is installed on the computer.

One criticism of using virtual machines to support computational reproducibility is that virtual-machine files are large (typically multiple gigabytes), especially if they include raw data files. This imposes a barrier for researchers to share virtual machines with the research community. One option is to use cloud-computing services (see Box 4). Scientists can execute an analysis in the cloud, take a “snapshot" of their virtual machine, and share it with others in that environment^95,98^. Cloud-based services typically provide repositories where virtual-machine files can be stored and shared easily among users. Despite these advantages, some researchers may prefer that their data reside on local computers, rather than in the cloud—at least while the research is being performed. In addition, cloud-based services may use proprietary software, so virtual machines may only be executable within each provider's infrastructure. Furthermore, to use a cloud-service provider, scientists may need to activate a fee-based account.

Another criticism of using virtual machines to support computational reproducibility is that the software and scripts used in the analysis will be less easily accessible to other scientists—details of the analysis are effectively concealed behind a “black box”^99^. Although other researchers may be able to reexecute the analysis within the virtual machine, it may be more difficult for them to understand and extend the analysis^99^. This problem can be ameliorated when all narratives, scripts, and code are stored in public repositories— separately from the virtual machine—and then imported when the analysis is executed^100^. Another solution is to use a prepackaged virtual machine, such as *Cloud BioLinux*, that contains a variety of software tools commonly used within a given research community^101^.

Scientists can automate the process of building and configuring virtual machines using tools such as *Vagrant* or *Vortex* (see Box 3). For either tool, users can write text-based configuration files that provide instructions for building virtual machines and allocating computational resources to them. In addition, these configuration files can be used to specify analysis steps^100^. Because these files are text based and relatively small (usually a few kilobytes), scientists can share them easily and track different versions of the files via source-control repositories. This approach also mitigates problems that might arise during the analysis stage. For example, even when a computer's *host* operating system must be reinstalled due to a computer hardware failure, the virtual machine can be recreated with relative ease.

#### Box 3 Virtual-machine software

Virtualization hypervisors:

- VirtualBox (open source) - https://www.virtualbox.org
- Xen (open source) - http://www.xenproject.org
- VMWare (partially open source) - http://www.vmware.com

Virtual-machine management tools:

- Vagrant (open source) - https://www.vagrantup.com
- Vortex (open source) - https://github.com/websecurify/node-vortex

#### Box 4 Commercial cloud-service providers

- Amazon Web Services - http://aws.amazon.com
- Rackspace Cloud - http://www.rackspace.com/cloud
- Google Cloud Platform - https://cloud.google.com/compute
- Windows Azure - https://azure.microsoft.com

## Software containers ease the process of installing and configuring dependencies

Software containers are a lighter-weight alternative to virtual machines. Like virtual machines, containers can encapsulate operating-system components, scripts, code, and data into a single package that can be shared with others. Thus, as with virtual machines, analyses executed within a software container should produce identical outputs, irrespective of the underlying operating system or whatever software may be installed outside the container (see Discussion for caveats). As is true for virtual machines, multiple containers can be executed simultaneously on a single computer, and each container may contain different software versions and configurations. However, whereas virtual machines include an entire operating system, software containers interface directly with the computer's main operating system and extend it as needed (Figure 3). This design provides less flexibility than virtual machines because containers are specific to a given type of operating system; however, containers require considerably less computational overhead than virtual machines and can be initialized much more quickly^102^.

The open-source *Docker* utility^103^—which has gained popularity among informaticians since its release in 2013—provides the ability to build, execute, and share software containers for Linux-based operating systems. Users specify a *Docker* container's contents using text-based commands. These instructions can be placed in a “Dockerfile," which other scientists can use to rebuild the container. As with virtual-machine configuration files, Dockerfiles are text based, so they can be shared easily and can be tracked and versioned in source-control repositories. Once a *Docker* container has been built, its contents can be exported to a binary file; these files are generally smaller than virtual-machine files, so they can be shared more easily—for example, via *DockerHub*^*104*^.

A key feature of *Docker* containers is that their contents can be stacked in distinct layers (or “images”). Each image includes software component(s) that address a particular need (see Figure 7 for an example). Within a given research lab, scientists might create general-purpose images that support functionality for multiple projects, and they might create specialized images that address the needs of specific projects. *Docker's* modular design provides the advantage that when images within a container are updated, *Docker* only needs to track the specific components that have changed; users who wish to update to a newer version must download a relatively small update. In contrast, even a minor change to a virtual machine would require users to export and reshare the entire virtual machine.

**Figure 7.**
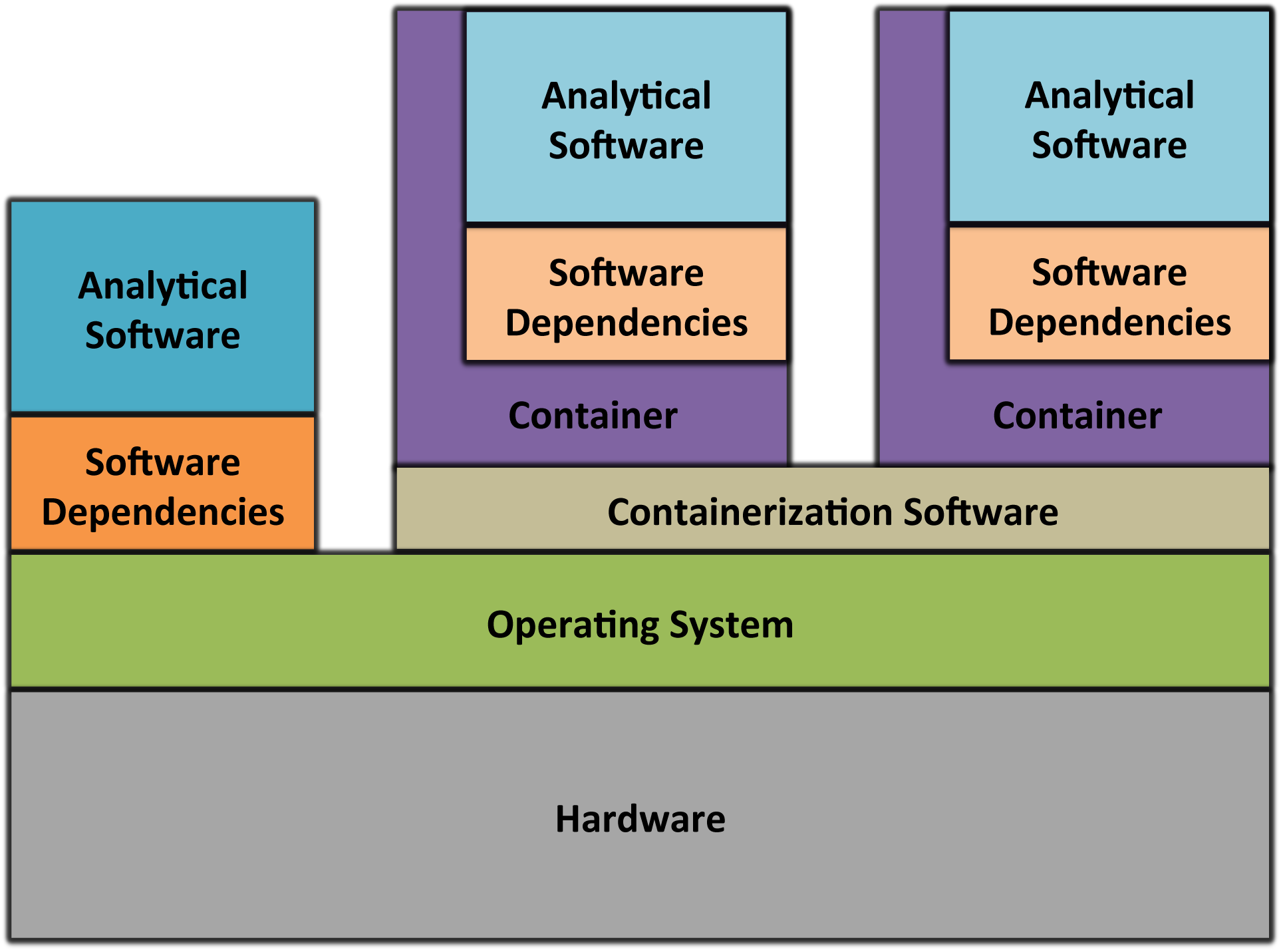
Architecture of software containers. Software containers encapsulate analytical software and dependencies. In contrast to virtual machines, containers execute within the context of the computer's main operating system.

Scientists have begun to share *Docker* images with others who are working in the same subdiscipline. For example, *nucleotid.es* is a catalog of genome-assembly tools that have been encapsulated in *Docker* images^105,106^. Genome-assembly tools differ considerably in the dependencies that they require and in the parameters that they support. This project provides a means to standardize these assemblers, to circumvent the need to install dependencies for each tool, and to perform benchmarks across the tools. Such projects may help to reduce the reproducibility burden on individual scientists.

The use of *Docker* containers for reproducible research comes with caveats. Individual containers are stored and executed in isolation from other containers on the same computer; however, because all containers on a given machine share the same operating system, this isolation is not as complete as it is with virtual machines. This means, for example, that a given container is not guaranteed to have access to a specific amount of computer memory or processing power—multiple containers may have to compete for these resources^102^. In addition, containers may be more vulnerable to security breaches^102^. Another caveat is that *Docker* containers can only be executed on Linux-based operating systems. For other operating systems, *Docker* containers must be executed within a virtual machine (for example, see http://boot2docker.io). Although this configuration offsets some benefits of using containers, combining virtual machines with containers may provide a happy medium for many scientists, allowing them to use a non-Linux *host* operating system, while receiving the benefits of containers within the *guest* operating system.

Efforts are ongoing to develop and refine software-container technologies. Box 5 lists various tools that are currently available. In coming years, these technologies promise to play an influential role within the scientific community.

#### Box 5 Open-source containerization software

- Docker - https://www.docker.com
- Linux Containers - https://linuxcontainers.org
- lmctfy - https://github.com/google/lmctfy
- OpenVZ - http://openvz.org
- Warden - http://docs.cloudfoundry.org/concepts/architecture/warden.html

## Discussion

Scientific advancement requires trust. This review provides a comprehensive, though inexhaustive, list of techniques that can help to engender such trust. Principally, scientists must perform research in such ways that they can trust their own findings^3,45^. Science philosopher Karl Popper contended that “[w]e do not take even our own observations quite seriously, or accept them as scientific observations, until we have repeated and tested them"^2^. Indeed, in many cases, the individuals who benefit most from computational reproducibility are those who performed the original analysis. But reproducible practices can also help scientists garner each other's trust^45,107^. When other scientists can reproduce an analysis and determine exactly how its conclusions were drawn, they may be more apt to cite the work and build upon it. In contrast, when others fail to reproduce research findings, it can derail scientific progress and lead to embarrassment, accusations, and retractions.

We have described seven tools and techniques for computational reproducibility. None of these approaches is sufficient for every scenario in isolation. Rather scientists will often find value in combining approaches. For example, a researcher who uses a literate-programming notebook (which combines narratives with code) might incorporate the notebook into a software container so that others can execute it without needing to install specific software dependencies. The container might also include a workflow-management system to ease the process of integrating multiple tools and incorporating best practices for the analysis. This container could be packaged within a virtual machine to ensure that it can be executed on many operating systems (see Figure 8). In determining a reproducibility strategy, scientists must evaluate the tradeoff between robustness and practicality.

**Figure 8.**
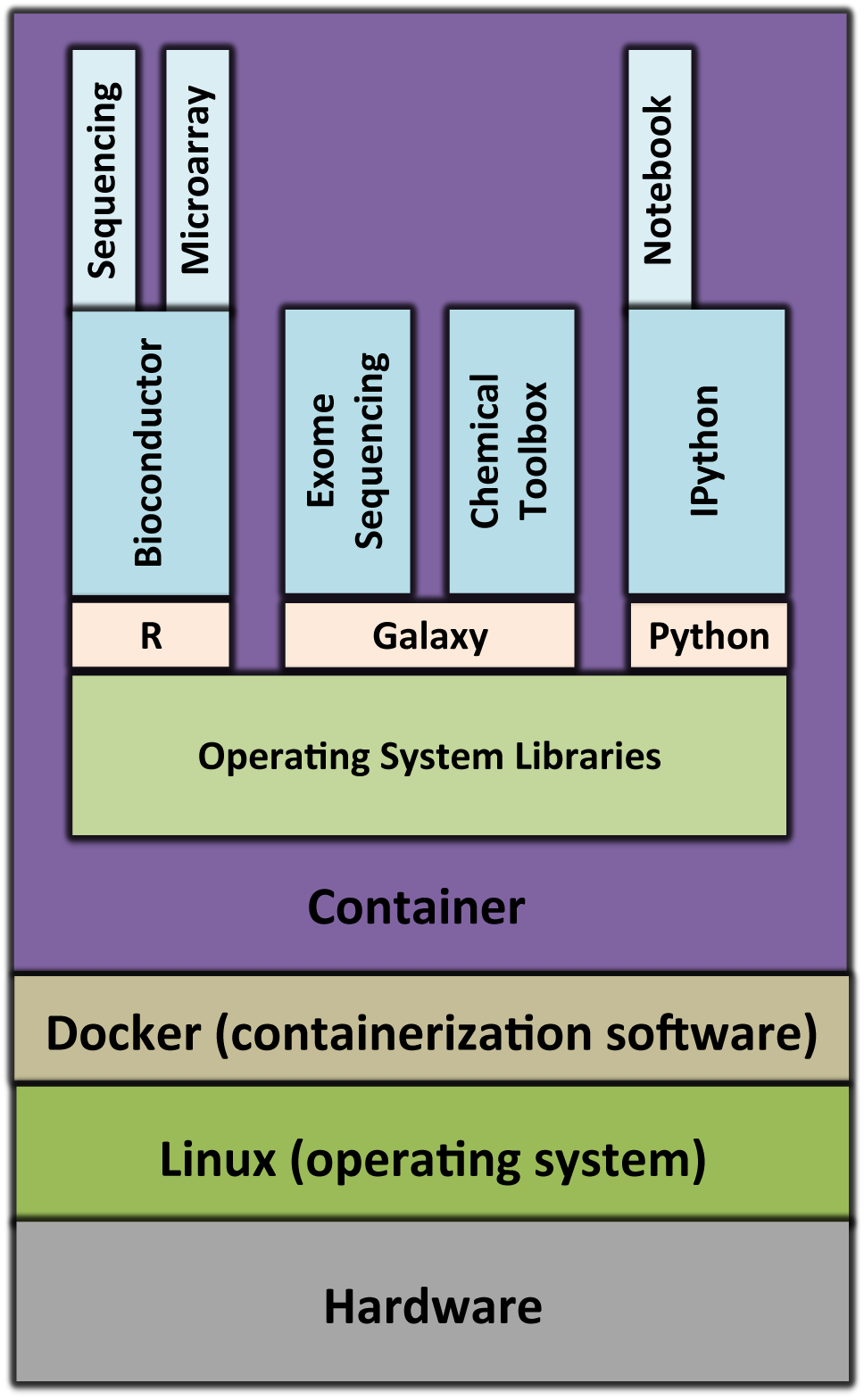
Example of a *Docker* container that could be used for genomics research. This container would enable researchers to preprocess various types of molecular data, using tools from *Bioconductor* and *Galaxy*, and to analyze the resulting data within an *IPython* notebook. Each box within the container represents a distinct *Docker* image. These images are layered such that some images depend on others (for example, the *Bioconductor* image depends on *R*). At its base, the container includes operating-system libraries, which may not be present (or may be configured differently) on the computer's main operating system.

The call for computational reproducibility relies on the premise that reproducible science will bolster the efficiency of the overall scientific enterprise^108^. Although reproducible practices may require additional time and effort, these practices provide ancillary benefits that help offset those expenditures^45^. Primarily, the scientists who perform a study may experience increased efficiency^45^. For example, before and after a manuscript is submitted for publication, it faces scrutiny from co-authors and peer reviewers who may suggest alterations to the analysis. Having a complete record of all analysis steps and being able to retrace those steps precisely, makes it faster and easier to implement the requested alterations^45,109^. Reproducible practices can also improve the efficiency of team science because colleagues can more easily communicate their research protocols and inspect each other's work; one type of relationship where this is critical is that between academic advisors and mentees^109^. Finally, when research protocols are shared transparently with the broader community, scientific advancement increases because scientists can learn more easily from each other's work and duplicate each other's efforts less frequently^109^.

Reproducible practices do not necessarily ensure that others can obtain results that are perfectly identical to what the original scientists obtained. Indeed, this objective may be infeasible for some types of computational analysis, including those that use randomization procedures, floating-point operations, or specialized computer hardware^96,110^. In such cases, the goal may shift to ensuring that others can obtain results that are semantically consistent with the original findings^5,6^. In addition, in studies where vast computational resources are needed to perform an analysis or where data sets are distributed geographically^111–113^, full reproducibility may be infeasible. Alternatively, it may be infeasible to reallocate computational resources for analyses that are highly computationally intensive^8^. In these cases, researchers can provide relatively simple examples that demonstrate the methodology^8^. When legal restrictions prevent researchers from sharing software or data publicly, or when software is available only via a Web interface, researchers should document the analysis steps as well as possible and describe why such components cannot be shared^24^.

Computational reproducibility does not guarantee against analytical biases or ensure that software produces scientifically valid results^114^. As with any research, a poor study design, confounding effects, or improper use of analytical software may plague even the most reproducible analyses^114,115^. On one hand, increased transparency puts scientists at a greater risk that such problems will be exposed. On the other hand, scientists who are fully transparent about their scientific approach may be more likely to avoid such pitfalls, knowing that they will be more vulnerable to such criticisms. Either way, the scientific community benefits.

Lastly, we emphasize that some reproducibility is better than none. As Voltaire said, the perfect should not be the enemy of the good^116^. Although some of the practices described in this review require more technical expertise than others, these practices are freely accessible to all scientists and provide long-term benefits to the researcher and to the scientific community. Indeed, as scientists act in good faith to perform these practices, where feasible, the pace of scientific progress will surely increase.

## References

1. Fisher, R. A. The Design of Experiments. (Hafner Press, 1935).

2. Popper, K. R. 2002. The logic of scientific discovery. (1959).

3. Peng, R. D. Reproducible research in computational science. Science 334, 1226–1227 (2011).

4. Russell, J. F. If a job is worth doing, it is worth doing twice. Nature 496, 7 (2013).

5. Feynman, R. P., Leighton, R. B. & Sands, M. Six Easy Pieces: Essentials of Physics Explained by Its Most Brilliant Teacher. 34–35 (Perseus Books, 1994).

6. Murray-Rust, P. & Murray-Rust, D. in Implementing Reproducible Research (eds. Stodden, V. C., Leisch, F. & Peng, R. D.) 113 (CRC Press, 2014).

7. Hey, A. J. G., Tansley, S., Tolle, K. M. & Others. The fourth paradigm: data-intensive scientific discovery. 1, (Microsoft Research Redmond, WA, 2009).

8. Millman, K. J. & Pérez, F. Developing Open-Source Scientific Practice. Implementing Reproducible Research 149 (2014).

9. Wilson, G. et al. Best practices for scientific computing. PLoS Biol. 12, e1001745 (2014).

10. Software with impact. Nat. Methods 11, 211 (2014).

11. Hong, N. C. We are the 92%. (2014). doi:10.6084/M9.FIGSHARE.1243288

12. Sacks, J., Welch, W. J., Mitchell, T. J. & Wynn, H. P. Design and Analysis of Computer Experiments. Stat. Sci. 4, 409–423 (1989).

13. Garijo, D. et al. Quantifying reproducibility in computational biology: The case of the tuberculosis drugome. PLoS One 8, (2013).

14. Error prone. Nature 487, 406 (2012).

15. Vandewalle, P., Barrenetxea, G., Jovanovic, I., Ridolfi, A. & Vetterli, M. Experiences with Reproducible Research in Various Facets of Signal Processing Research. in 2007 IEEE International Conference on Acoustics, Speech and Signal Processing - ICASSP’07 4, IV-1253-IV-1256 (IEEE, 2007).

16. Cassey, P., Cassey, P., Blackburn, T. & Blackburn, T. Reproducibility and Repeatability in Ecology. Bioscience 56, 958–959 (2006).

17. Murphy, J. M. et al. Quantification of modelling uncertainties in a large ensemble of climate change simulations. Nature 430, 768–772 (2004).

18. McCarthy, D. J. et al. Choice of transcripts and software has a large effect on variant annotation. Genome Med. 6, 26 (2014).

19. Neuman, J. A., Isakov, O. & Shomron, N. Analysis of insertion-deletion from deep-sequencing data: Software evaluation for optimal detection. Brief. Bioinform. 14, 46–55 (2013).

20. Bradnam, K. R. et al. Assemblathon 2: evaluating de novo methods of genome assembly in three vertebrate species. Gigascience 2, 10 (2013).

21. Bilal, E. et al. Improving Breast Cancer Survival Analysis through Competition-Based Multidimensional Modeling. PLoS Comput. Biol. 9, e1003047 (2013).

22. Moskvin, O. V., McIlwain, S. & Ong, I. M. CAMDA 2014: Making sense of RNA-Seq data: From low-level processing to functional analysis. Systems Biomedicine 2, 31–40 (2014).

23. Reducing our irreproducibility. Nature 496, 398–398 (2013).

24. Evolution of Translational Omics: Lessons Learned and the Path Forward. (The National Academies Press, 2012).

25. Collins, F. S. & Tabak, L. a. Policy: NIH plans to enhance reproducibility. Nature 505, 612–613 (2014).

26. Chambers, J. M. S as a Programming Environment for Data Analysis and Graphics. in Problem Solving Environments for Scientific Computing, Proc. 17th Symp. on the Interface of Stat. and Comp. 211–214 (1985).

27. LeVeque, R. J., Mitchell, I. M. & Stodden, V. Reproducible research for scientific computing: Tools and strategies for changing the culture. Computing in Science and Engineering 14, 13 (2012).

28. Stodden, V., Guo, P. & Ma, Z. Toward Reproducible Computational Research: An Empirical Analysis of Data and Code Policy Adoption by Journals. PLoS One 8, 2–9 (2013).

29. Morin, A. et al. Research priorities. Shining light into black boxes. Science 336, 159–160 (2012).

30. Rebooting review. Nat. Biotechnol. 33, 319 (2015).

31. Ioannidis, J. P. a. et al. Repeatability of published microarray gene expression analyses. Nat. Genet. 41, 149–155 (2009).

32. Nekrutenko, A. & Taylor, J. Next-generation sequencing data interpretation: enhancing reproducibility and accessibility. Nat. Rev. Genet. 13, 667–672 (2012).

33. Baggerly, K. a. & Coombes, K. R. Deriving chemosensitivity from cell lines: Forensic bioinformatics and reproducible research in high-throughput biology. Ann. Appl. Stat. 3, 1309–1334 (2009).

34. Decullier, E., Huot, L., Samson, G. & Maisonneuve, H. Visibility of retractions: a cross-sectional one-year study. BMC Res. Notes 6, 238 (2013).

35. Claerbout, J. F. & Karrenbach, M. Electronic Documents Give Reproducible Research a New Meaning. in Meeting of the Society of Exploration Geophysics (1992).

36. Stodden, V. & Miguez, S. Best Practices for Computational Science: Software Infrastructure and Environments for Reproducible and Extensible Research. Journal of Open Research Software 2, 21 (2014).

37. Ravel, J. & Wommack, K. E. All hail reproducibility in microbiome research. Microbiome 2, 8 (2014).

38. Stodden, V. 2014: What scientific idea is ready for retirement? *http://edge.org/response-detail/25340* (2014). at <http://edge.org/response-detail/25340>

39. Birney, E. et al. Prepublication data sharing. Nature 461, 168–170 (2009).

40. Hothorn, T. & Leisch, F. Case studies in reproducibility. Brief. Bioinform. 12, 288–300 (2011).

41. Schofield, P. N. et al. Post-publication sharing of data and tools. Nature 461, 171–173 (2009).

42. Piwowar, H. a., Day, R. S. & Fridsma, D. B. Sharing detailed research data is associated with increased citation rate. PLoS One 2, (2007).

43. Johnson, V. E. Revised standards for statistical evidence. Proc. Natl. Acad. Sci. U. S. A. 110, 19313–19317 (2013).

44. Halsey, L. G., Curran-everett, D.., Vowler, S. L. & Drummond, G. B. The fickle P value generates irreproducible results. Nat. Methods 12, 179–185 (2015).

45. Sandve, G. K., Nekrutenko, A., Taylor, J. & Hovig, E. Ten Simple Rules for Reproducible Computational Research. PLoS Comput. Biol. 9, 1–4 (2013).

46. Foundation, F. S. GNU Make at <https://www.gnu.org/software/make>

47. Make for Windows. at <http://gnuwin32.sourceforge.net/packages/make.htm>

48. Koster, J. & Rahmann, S. S nakemake—a scalable bioinformatics workflow engine. Bioinformatics 28, 2520–2522 (2012).

49. Sadedin, S. P., Pope, B. & Oshlack, A. Bpipe: A Tool for Running and Managing Bioinformatics Pipelines. Bioinformatics 28, 1525–1526 (2012).

50. Tange, O. GNU Parallel - The Command-Line Power Tool. ;login: The USENIXMagazine 36, 4247 (2011).

51. Albrecht, M., Donnelly, P., Bui, P. & Thain, D. Makeflow: A portable abstraction for data intensive computing on clusters, clouds, and grids. in Proceedings of the 1st ACM SIGMOD Workshop on Scalable Workflow Execution Engines and Technologies (2012).

52. Knight, S., Austin, C., Crain, C., Leblanc, S. & Roach, A. Scons software construction tool. (2011). at <http://www.scons.org>

53. CMake. at <https://cmake.org>

54. Code share. Nature 514, 536 (2014).

55. Loeliger, J. & McCullough, M. Version Control with Git: Powerful Tools and Techniques for Collaborative Software Development. 456 (‘O’Reilly Media, Inc.’, 2012).

56. Huber, W. et al. Orchestrating high-throughput genomic analysis with Bioconductor. Nat. Methods 12, 115–121 (2015).

57. R Core Team. R: A Language and Environment for Statistical Computing. (2014). at <http://www.r-project.org/>

58. Tóth, G. et al. Space Weather Modeling Framework: A new tool for the space science community. J. Geophys. Res. 110, A12226 (2005).

59. Tan, E., Choi, E., Thoutireddy, P., Gurnis, M. & Aivazis, M. GeoFramework: Coupling multiple models of mantle convection within a computational framework. Geochem. Geophys. Geosyst. 7, (2006).

60. Heisen, B. et al. Karabo: An Integrated Software Framework Combining Control, Data Management, and Scientific Computing Tasks. in 14th International Conference on Accelerator & Large Experimental Physics Control Systems, ICALEPCS2013 (2013).

61. Schneider, C. A., Rasband, W. S. & Eliceiri, K. W. NIH Image to ImageJ: 25 years of image analysis. Nat. Methods 9, 671–675 (2012).

62. Schindelin, J. et al. Fiji: an open-source platform for biological-image analysis. Nat. Methods 9, 676–682 (2012).

63. Biasini, M. et al. OpenStructure: an integrated software framework for computational structural biology. Acta Crystallogr. D Biol. Crystallogr. 69, 701–709 (2013).

64. Home | Apache Ivy. at <http://ant.apache.org/ivy/>

65. Home. Puppet Labs at <https://puppetlabs.com>

66. Martin, R. C. Clean code: a handbook of agile software craftsmanship. (Pearson Education, 2009).

67. Knuth, D. E. Literate Programming. Comput. J. 27, 97–111 (1984).

68. Pérez, F. & Granger, B. E. IPython: a System for Interactive Scientific Computing. Computing in Science and Engineering 9, 21–29 (2007).

69. Shen, H. Interactive notebooks: Sharing the code. Nature 515, 151–152 (2014).

70. Project Jupyter. at <https://jupyter.org>

71. Xie, Y. Dynamic Documents with R and knitr. 216 (CRC Press, 2013).

72. RStudio Team. RStudio: Integrated Development for R. at <http://www.rstudio.com>

73. Gross, A. M. et al. Multi-tiered genomic analysis of head and neck cancer ties TP53 mutation to 3p loss. Nat. Genet. 46, 1–7 (2014).

74. Ding, T. & Schloss, P. D. Dynamics and associations of microbial community types across the human body. Nature 509, 357–360 (2014).

75. Ram, Y. & Hadany, L. The probability of improvement in Fisher's geometric model: A probabilistic approach. Theor. Popul. Biol. 99, 1–6 (2015).

76. Meadow, J. F. et al. Bacterial communities on classroom surfaces vary with human contact. Microbiome 2, 7 (2014).

77. White, E. Programming for Biologists. at <http://www.programmingforbiologists.org>

78. Wilson, G. Software Carpentry: lessons learned. F1000Res. 3, 62 (2014).

79. Peng, R. D. Coursera course: Computing for Data Analysis. at <https://www.coursera.org/course/compdata>

80. Bioconductor - Courses and Conferences. at <http://master.bioconductor.org/help/course-materials>

81. Gil, Y. et al. Examining the challenges of scientific workflows. Computer 40, 24–32 (2007).

82. Giardine, B. et al. Galaxy: a platform for interactive large-scale genome analysis. Genome Res. 15, 1451–1455 (2005).

83. Goecks, J., Nekrutenko, A. & Taylor, J. Galaxy: a comprehensive approach for supporting accessible, reproducible, and transparent computational research in the life sciences. Genome Biol. 11, R86 (2010).

84. Afgan, E. et al. Harnessing cloud computing with Galaxy Cloud. Nat. Biotechnol. 29, 972–974 (2011).

85. Galaxy. at <https://usegalaxy.org>

86. Callahan, S. P. et al. VisTrails: Visualization Meets Data Management. in Proceedings of the 2006 ACM SIGMOD International Conference on Management of Data 745–747 (ACM, 2006).

87. Davidson, S. B. & Freire, J. Provenance and scientific workflows. in Proceedings of the 2008 ACM SIGMOD international conference on Management of data - SIGMOD’08 1345 (2008).

88. Altintas, I. et al. Kepler: an extensible system for design and execution of scientific workflows. in Proceedings. 16th International Conference on Scientific and Statistical Database Management, 2004. 423–424 (IEEE, 2004).

89. Goff, S. A. et al. The iPlant Collaborative: Cyberinfrastructure for Plant Biology. Front. Plant Sci. 2, 34 (2011).

90. Reich, M. et al. GenePattern 2.0. Nat. Genet. 38, 500–501 (2006).

91. Reich, M. et al. GenomeSpace: An environment for frictionless bioinformatics. Cancer Res. 72, 3966–3966 (2012).

92. Wolstencroft, K. et al. The Taverna workflow suite: designing and executing workflows of Web Services on the desktop, web or in the cloud. Nucleic Acids Res. 41, 557–561 (2013).

93. Rex, D. E., Ma, J. Q. & Toga, A. W. The LONI Pipeline Processing Environment. Neuroimage 19, 1033–1048 (2003).

94. Lazarus, R., Kaspi, A. & Ziemann, M. Creating re-usable tools from scripts: The Galaxy Tool Factory. Bioinformatics 28, 3139–3140 (2012).

95. Dudley, J. T. & Butte, A. J. In silico research in the era of cloud computing. Nat. Biotechnol. 28, 1181–1185 (2010).

96. Hurley, D. G., Budden, D. M. & Crampin, E. J. Virtual Reference Environments: a simple way to make research reproducible. Brief. Bioinform. 1–3 (2014).

97. Gent, I. P. The Recomputation Manifesto. arXiv (2013). at <http://arxiv.org/abs/1304.3674>

98. Howe, B. Virtual Appliances, Cloud Computing, and Reproducible Research. Comput. Sci. Eng. 14, 36–41 (2012).

99. Brown, C. T. Virtual machines considered harmful for reproducibility. (2012). at <http://ivory.idyll.org/blog/vms-considered-harmful.html>

100. Piccolo, S. R. Building portable analytical environments to improve sustainability of computational-analysis pipelines in the sciences. (2014). doi:10.6084/m9.figshare.1112571

101. Krampis, K. et al. Cloud BioLinux: pre-configured and on-demand bioinformatics computing for the genomics community. BMC Bioinformatics 13, 42 (2012).

102. Felter, W., Ferreira, A., Rajamony, R. & Rubio, J. An Updated Performance Comparison of Virtual Machines and Linux Containers. (2014). at <http://domino.research.ibm.com/library/CyberDig.nsf/papers/0929052195DD819C85257D2300681e7B/$File/rc25482.pdf>

103. docker. docker at <https://www.docker.com>

104. Docker Hub. at <https://hub.docker.com>

105. Belmann, P. et al. Bioboxes: standardised containers for interchangeable bioinformatics software. Gigascience 4, 47 (2015).

106. Barton, M. nucleotides · genome assembler benchmarking. at <http://nucleotid.es>

107. Hones, M. J. Reproducibility as a Methodological Imperative in Experimental Research. in PSA: Proceedings of the Biennial Meeting of the Philosophy of Science Association One, 585–599 (Philosophy of Science Association, 1990).

108. Crick, T. ‘Share and Enjoy’: Publishing Useful and Usable Scientific Models. at <http://arxiv.org/abs/1409.0367v2>

109. Donoho, D. L. An invitation to reproducible computational research. Biostatistics 11, 385–388 (2010).

110. Goldberg, D. What Every Computer Scientist Should Know About Floating-point Arithmetic. ACM Comput. Surv. 23, 5–48 (1991).

111. Shirts, M. & Pande, V. S. COMPUTING: Screen Savers of the World Unite! Science 290, 1903–1904 (2000).

112. Bird, I. Computing for the Large Hadron Collider. Annu. Rev. Nucl. Part. Sci. 61, 99–118 (2011).

113. Anderson, D. P. BOINC: A System for Public Resource Computing and Storage. in Proceedings of the Fifth IEEE/ACM International Workshop on Grid Computing (GRID’04) (2004).

114. Ransohoff, D. F. Bias as a threat to the validity of cancer molecular-marker research. Nat. Rev. Cancer 5, 142–149 (2005).

115. Bild, A. H., Chang, J. T., Johnson, W. E. & Piccolo, S. R. A field guide to genomics research. PLoS Biol. 12, e1001744 (2014).

116. Ratcliffe, S. Concise Oxford Dictionary of Quotations. 389 (Oxford University Press, 2011).

